# Integrated single-cell analysis reveals interferon-driven immune signatures in a DENV1 human infection model

**DOI:** 10.64898/2026.01.04.697597

**Authors:** A.S. Vermeersch, T.W. Verdonckt, C. Struyfs, E. De Meester, A. Waickman, S.J. Thomas, K.K. Ariën, O. Lagatie, F. Van Nieuwerburgh

## Abstract

Dengue virus infection triggers complex innate and adaptive immune responses, yet the molecular mechanisms that shape early antiviral immunity remain incompletely defined. We performed longitudinal single-cell multi-omics profiling of peripheral blood mononuclear cells from four flavivirus-naïve adults experimentally infected with DENV-1, integrating 5′ scRNA-seq with paired surface proteomics.

Across 95,841 high-quality cells collected at baseline and on days 8 and 10 post-infection, we observed a strong interferon-driven transcriptional response accompanied by marked immune-cell redistribution, including expansion of monocytes and transient reductions in dendritic cells and double-negative T cells. Cytotoxic and helper lymphocyte populations, particularly naïve, central memory, and effector memory CD8⁺ T cells, showed extensive crosstalk with monocytes and NK cells, reflecting coordinated cytokine production and cytotoxic activation. Early B cell activation was evident through increased immunoglobulin gene expression. Innate sensing pathways, including RIG-I and Toll-like signaling, were activated across NK, T, and B cell subsets, while also demonstrating enrichment of antigen processing and apoptosis programs. Pro-inflammatory and cytotoxic signatures peaked at day 8, supported by broad upregulation of interferon-stimulated, pro-apoptotic, and regulatory genes.

Together, these findings define a robust IFN-driven antiviral state and coordinated activation of immune cell subsets, providing new insights into the immune dynamics of primary dengue infection.

**Importance:** Dengue virus infects millions of people each year, but the early immune events that shape disease outcomes are still unclear. Most studies measure average responses across all blood cells, which hides how individual cell types react. By tracking thousands of single immune cells from volunteers infected with dengue virus under controlled conditions, we show how the immune system rapidly reorganizes during the first days of infection. Many cell types activate antiviral programs, communicate with one another, and shift their behavior in a coordinated way, driven by strong interferon activity. These results provide a clearer view of how early immune responses unfold in humans and identify cellular processes that may influence who develops more severe illness.

## Introduction

Dengue is a mosquito-borne viral disease caused by four antigenically distinct DENV serotypes and remains a major cause of illness in tropical and subtropical regions (1). Symptomatic illness is classified as dengue without warning signs (DWoWS), dengue with warning signs (DWWS), or severe dengue (SD). Most individuals experience self-limiting dengue, whereas approximately 20% develop warning signs such as abdominal pain, persistent vomiting, or mucosal bleeding (2). Severe dengue occurs in roughly 5% of symptomatic cases, often during defervescence, and can lead to hemorrhagic fever or shock (2, 3). Disease progression is shaped both by viral pathogenesis and the magnitude of the host inflammatory response (4, 5). Although antibody-dependent enhancement contributes to severe disease during heterologous secondary infection, severe outcomes also occur in primary infections, indicating additional mechanisms of immune dysregulation (5–7). Cross-reactivity to related flaviviruses, including Zika virus, may further modulate disease risk (8).

Interferon-driven antiviral programs and inflammatory chemokines are central features of acute dengue, with monocytes and dendritic cells serving as major targets of infection and key coordinators of innate immunity (9–13). T and B cell responses also influence disease outcome, yet the cellular interactions that shape these responses *in vivo* remain incompletely defined (14–17). Most transcriptomic studies rely on bulk profiling, obscuring cellular heterogeneity and limiting inference on intercellular communication. Recent single-cell studies have begun to address these limitations but have focused largely on transcriptomes without incorporating protein-level phenotypes or modeling coordinated signaling between immune compartments (13, 18, 19).

Human challenge models have successfully reproduced key clinical, serological, and virological features of natural primary DENV infections under controlled conditions (20–22). Waickman et al. (2021) demonstrated that experimental infection recapitulates core transcriptional features of natural primary disease. However, their analyses emphasized V(D)J repertoires and gene expression, and did not integrate multimodal single-cell measurements or intercellular network modeling (20). As a result, the dynamic coordination of innate and adaptive immune subsets during early DENV-1 infection remains incompletely resolved.

Here, we extend prior work by combining 5′ scRNA-seq with surface proteomics to profile PBMCs from flavivirus-naïve adults experimentally infected with DENV-1. Integrating multimodal data with intercellular communication and pathway analyses, we identify monocyte-centered rewiring of intercellular signaling networks, coordinated remodeling of lymphocyte activation programs, early B-cell adhesion and activation programs across CD4⁺, CD8⁺, NK, and γδ T cell subsets, and widespread suppression of translational pathways across lymphoid compartments. Together, these findings define a coordinated, multimodal immune response during primary dengue infection and establish a foundation for mechanistic insight and biomarker discovery.

## Material & methods

### Study design

The DENV-1 human infection model was conducted as an open-label Phase 1 study (ClinicalTrials.gov NCT03869060) (20). Nine flavivirus-naïve adults were inoculated subcutaneously with the underattenuated DENV-1 strain 45AZ5, and PBMCs were collected longitudinally. From this cohort, we analyzed twelve PBMC samples obtained from four participants on days 0, 8, and 10 post-infection, corresponding to pre-infection and the febrile phase. All participants provided written informed consent, and the study was approved by the appropriate institutional review boards.

### Single-cell multi-omic profiling

Cryopreserved PBMCs were processed using the 10X Genomics 5′ single-cell multi-omics workflow with paired gene expression and surface-protein (TotalSeq-C) libraries. Libraries were prepared according to the manufacturer’s protocol and sequenced to a target depth of minimum 20,000 paired-end reads per cell for gene expression (GEX) and 5,000-10,000 reads per cell for surface proteins (FB).

### Data processing and quality control

Sequencing data were processed with Cell Ranger with GRCh38 as reference genome. Filtered gene expression and surface protein matrices were analyzed in R using Seurat. Standard quality-control procedures were applied to remove low-quality cells and doublets, and datasets were integrated using Harmony. Weighted nearest-neighbor analysis was used for multimodal clustering and annotation.

### Differential abundance and gene expression analysis

Cell-type abundance changes were quantified using negative-binomial generalized linear models implemented in edgeR. Pseudobulk differential gene expression analyses were performed with muscat. Differential protein abundance was assessed using Seurat’s FindMarkers function in a paired design. Significance thresholds were set at adjusted p-value < 0.05 and |log₂ fold change| ≥ 1. Gene ontology (GO) and Kyoto Encyclopedia of Genes and Genomes (KEGG) enrichment analysis for marker genes of cell clusters was done using fgsea and msigdbr.

### Intercellular communication analysis

Ligand–receptor interactions were inferred using CellChat, applied separately to day 0, day 8, and day 10 datasets. Pathway-level signaling probabilities were computed using aggregated ligand–receptor pairs within each pathway.

### Viral RNA detection

DENV-1 RNA reads were quantified using Cell Ranger and Vulture with a combined human-viral reference. Viral transcripts were retained after filtering host-derived UMIs.

Methodological details, parameter settings, and software versions are provided in **Supplementary File S1**.

## Results

### Clinical characteristics

All participants developed detectable levels of viral RNA in their serum (**Figure 1**) (20). Peak viral loads were observed on day 8-9 for participants 204 and 205, while a delayed peak was seen in participants 208 and 209 on day 11. Due to the limited sample size, further stratification of the groups based on the timing of their viral load peak was not feasible, as the resulting subgroup sizes would lack statistical power. All four individuals experienced solicited systemic adverse events following inoculation, including headache, fatigue or weakness, eye pain, myalgia, rash (4/4), leukopenia (3/4), fever (3/4), and arthralgia (2/4) (**Supplementary Table S1**). Three participants required hospitalization: participant 204 for three days, and participants 208 and 209 for seven days.

**Figure 1.**
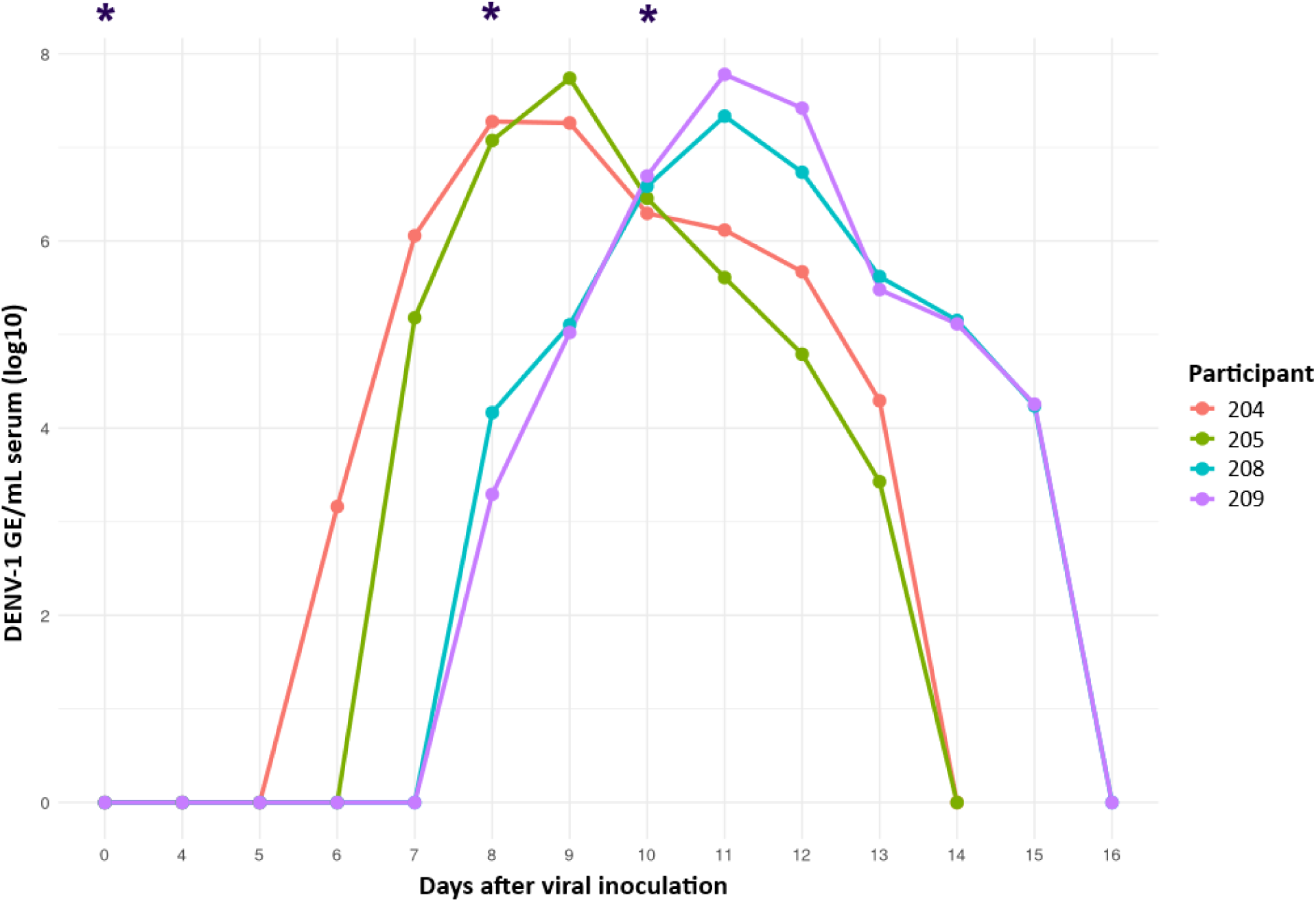
Serum viral load trajectories for four participants from day 0 to day 16, measured by quantitative real-time PCR as reported in Waickman et al. (2021)(**20**). Blue asterisks mark the time points analyzed in the present study.

### Profile of the human PBMCs at single-cell resolution

The mean sequencing depth was 31,985 reads per cell (SD = 11,665.5) with an average of 299,750,755 total reads per sample and an average of 152,095 cells captured before filtering the GEX data. The FB data had an average of 13,723.5 reads per cell (SD = 7,948). After quality control 95,841 cells were retained for downstream analysis.

Following clustering based on gene expression, cells were visualized using weighted nearest neighbor analysis using uniform manifold approximation and projection for dimension reduction (wnnUMAP,), and annotated with Azimuth based on the expression of canonical genes (23) (**Supplementary Table S2)**. The final dataset contained eight major cell clusters: B cells, CD4^+^ T cells, CD8^+^ T cells, dendritic cells (DC), monocytes, natural killer cells (NK), ‘other T cells’ such as γδ T cells, mucosal-associated invariant T cells (MAIT), double-negative T cells (dnT), and a group labeled as “other” containing non-lymphoid hematopoietic lineages such as innate lymphoid cells (ILC), erythroid cells, platelets and hematopoietic stem and progenitor cells (HSPC).

Further subclassification revealed 26 immune cell subtypes (**Figure 2A**), including naïve, central memory (TCM), effector memory (TEM), and cytotoxic subsets of both CD4⁺ and CD8⁺ T cells, and proliferating CD8⁺ T cells; γδ T cells; dnT; regulatory T cells (Tregs); and MAIT. Among B cells, naïve, intermediate, memory B cells, and plasmablasts were distinguished. The monocyte population consisted of CD14⁺ and CD16⁺ subsets, while the dendritic cell (DC) compartment included plasmacytoid DCs (pDCs). Additionally, NK cells were further divided into canonical, proliferating, and CD56bright subsets, and a separate cluster of ILCs was identified. Rare cell populations included hematopoietic stem and progenitor cells (HSPCs), erythroid cells, and platelets.

**Figure 2.**
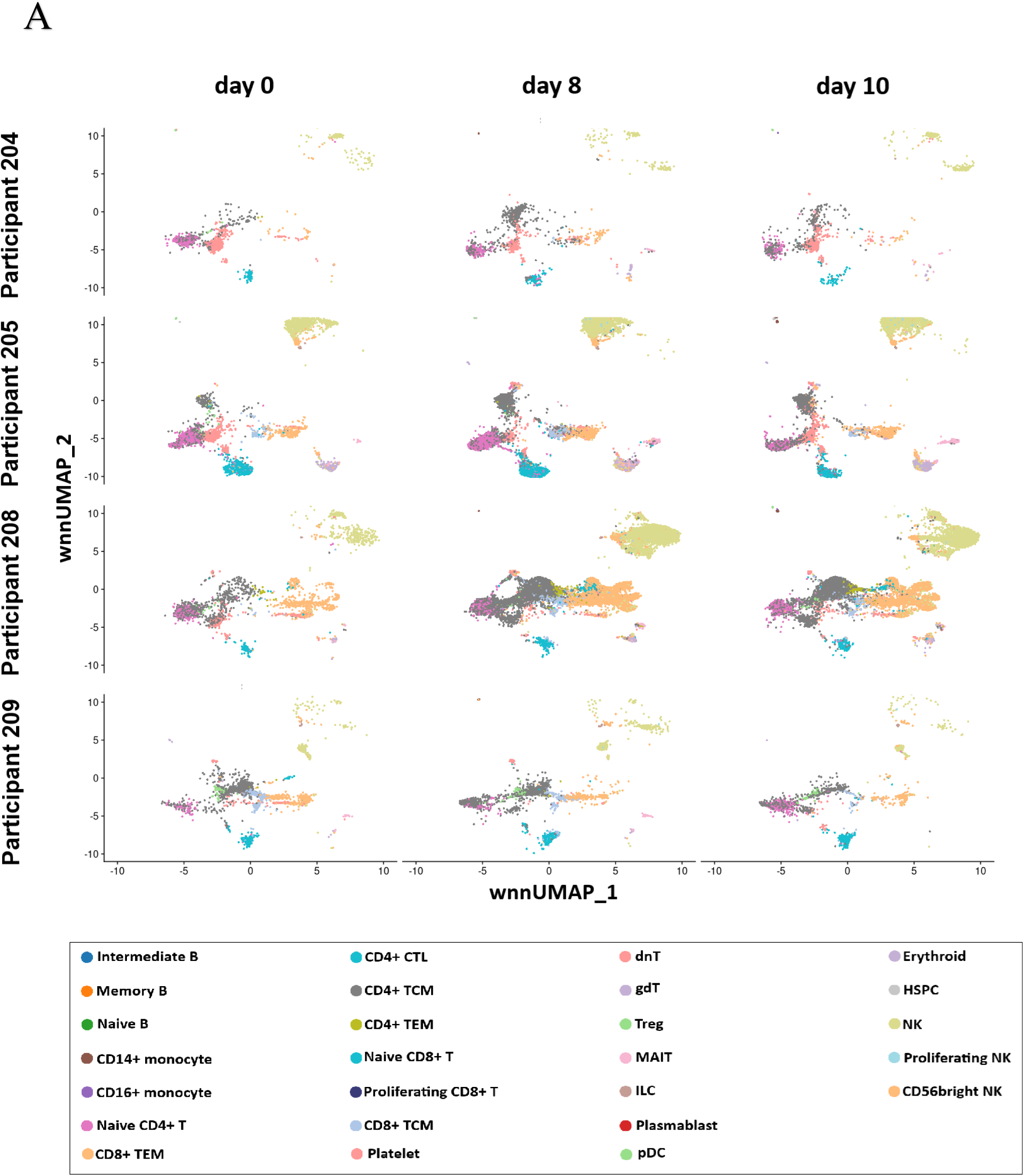
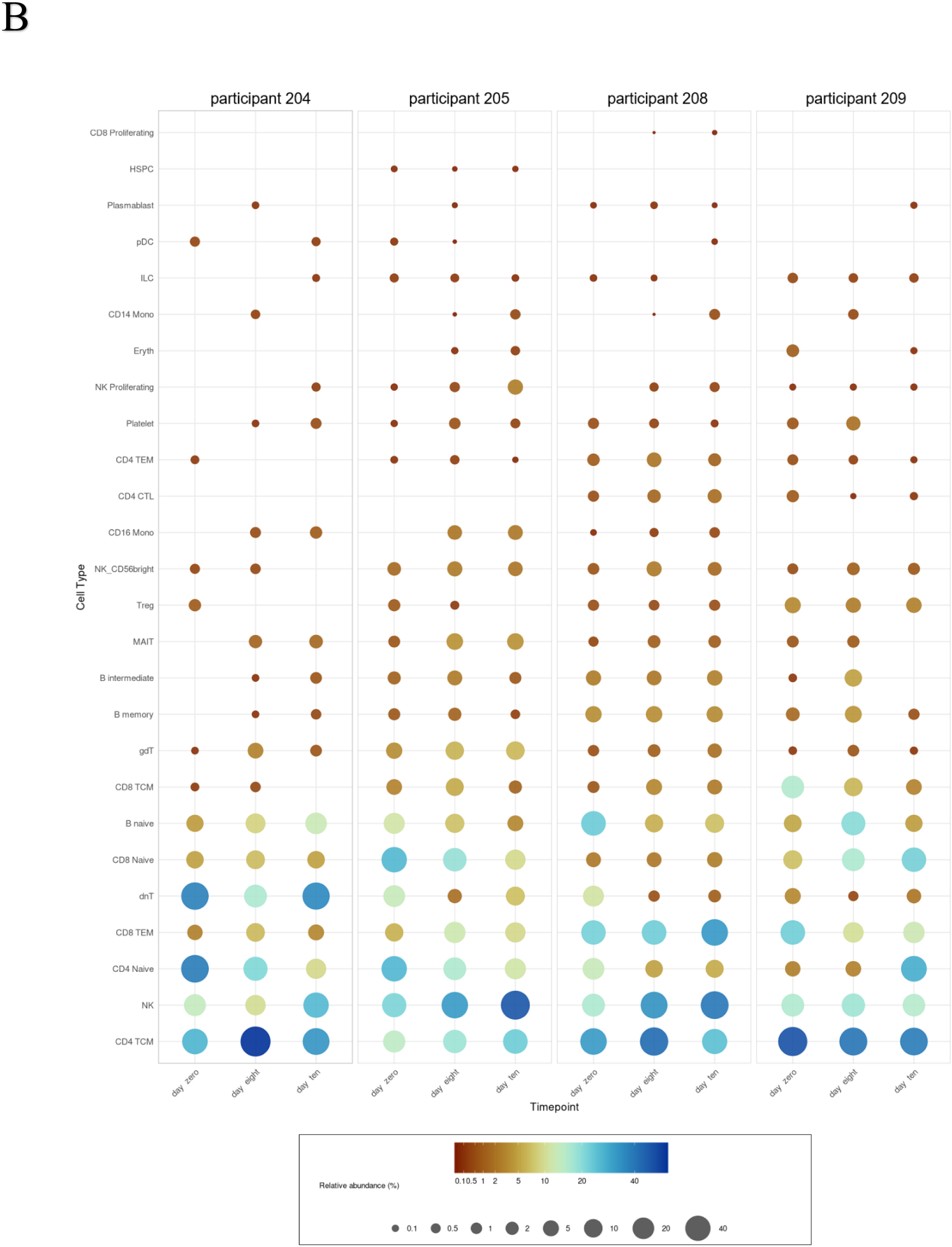
(A) wnnUMAP projections of major immune cell clusters for each participant at days 0, 8, and 10 post infection. cDC2 are not shown due to low abundance. (B) Relative abundances of immune cell subtypes across all time points, displayed separately for each participant. Abbreviations: FDR, false discovery rate; wnnUMAP, weighted nearest neighbor analysis using uniform manifold approximation and projection for dimension reduction.

Absolute cell counts (**Supplementary Figure S1**) and relative proportions (**Figure 2B**) of each major cell subtype were calculated for each participant and time point.

Monocyte abundance significantly increased at D8 (logFC = 6.370; padj = 0.005142) and remained elevated at D10 (logFC = 6.70; padj =0.00267) relative to baseline (D0). At the subtype level, dnT cells exhibited a marked decrease in abundance from D0 to D8 (logFC = –3.21; padj = 0.01578). When we applied an exploratory stratification by timing of the viral load peak (early vs late) using an unpaired design to avoid collinearity, slightly different patterns emerged. In participants with an early viral load peak, monocyte abundance increased significantly from D0 to D8/D10 (logFC = 9.40 and 9.75, padj = 0.001 and 0.0017, respectively), whereas no significant changes were observed in the late peak group (logFC = 3.72 and 4.45, padj = 0.290 and 0.094, respectively).

At the subtype level, CD16⁺ monocytes expanded markedly yet not significantly by D8/D10 in early peak participants (logFC = 9.25 and 9.57; padj = 0.086 and 0.003, respectively), consistent with early myeloid activation. Interaction terms between viral load group and time point were not significant, suggesting comparable temporal trajectories across groups.

### Differential gene expression analysis

Differentially expressed genes (DEGs) were identified among cell subtype clusters (**Figure 3; Supplementary File S2**), and the top ten genes per cell subtype are listed in **Supplementary Tables S3a–e** (|logFC| ≥ 1; padj < 0.05). The overlap between time point comparisons within cell subtype clusters is shown in **Supplementary Figure S2**. In total, 979 genes over 9 different cell clusters were upregulated in D8 compared to D0, 200 genes when comparing D10 to D0 over 9 cell clusters and just 1 gene (HBB) of γδ T cells was upregulated on D10 compared to D8. On D8 compared to D0, 642 genes were downregulated over 9 cell types and 41 genes in the comparison D10 to D0 over 7 distinct cell types. On D10, 6 genes over 5 cell subtypes were downregulated compared to D8. Several cell types were excluded from the muscat analysis due to not passing the minimum cell count threshold of 10 cells per sample – cell type combination. These included “other cells”, monocytes and DCs, and on a deeper level intermediate B cells, B memory cells, CD14^+^ monocytes, CD16^+^ monocytes, CD4^+^ CTL, CD4^+^ TEM, proliferating CD8^+^ T cells, CD8^+^ TCM, erythroid cells, HSPC, ILC, proliferating NK cells, NK CD56bright cells, pDC, plasmablasts and platelets.

**Figure 3.**
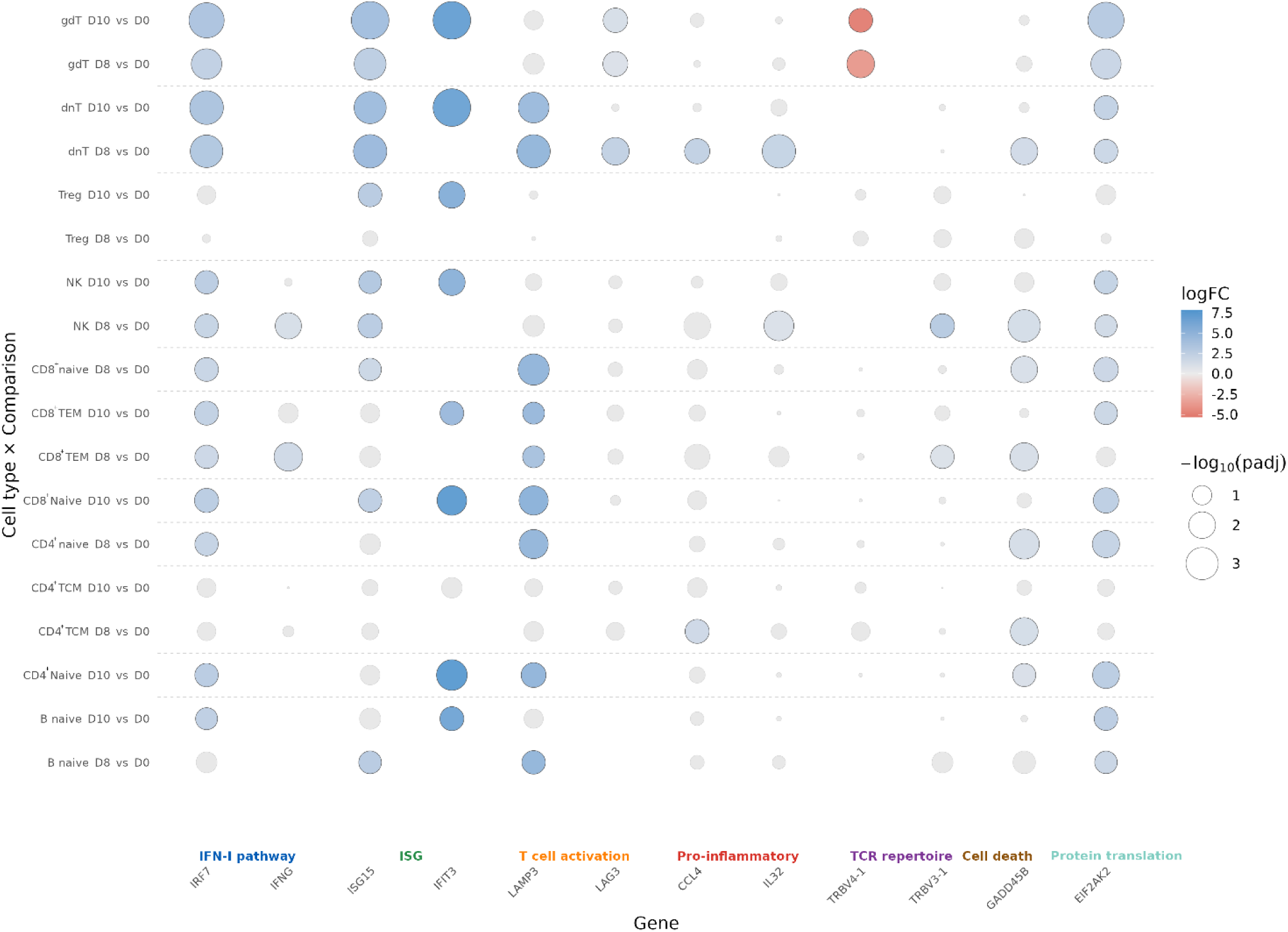
Dot plot showing selected results from differential gene expression analyses for the D0–D8 and D0–D10 comparisons, performed using muscat. Results were filtered for adjusted p value (padj) < 0.05 and absolute log₂ fold change |logFC| ≥ 1. Genes were selected based on previously reported relevance in the literature (6, 12, 13, 18)**. Each dot represents a gene-cell type pair, with dot size proportional to statistical significance (−log₁₀ padj) and color indicating the direction and magnitude of expression change (logFC).**

Interferon-stimulated genes (ISGs) dominated the transcriptional landscape on both D8 and D10. Canonical ISGs, including IFI44L, ISG15, MX2, OAS1, GBP1, USP18, and members of the IFIT family, were robustly upregulated across multiple cell types. This was accompanied by increased expression of key IFN signaling mediators, such as STAT1 and IRF7, as well as the type II interferon IFNγ, indicating activation of both type I and type II IFN pathways and a robust systemic antiviral response.

Translational control pathways were dynamically regulated, reflecting coordinated modulation of the host translation machinery during infection. EIF4E3, involved in translation initiation and elongation, was downregulated in CD8⁺ TEM, and γδ T cells on D8. In contrast, EIF2AK2 was upregulated in NK cells, γδ T cells, dnT cells, naïve B cells, and CD4⁺ and CD8⁺ T cells on D8, expanded to CD8⁺ TEM on D10. EIF5A was similarly upregulated in NK cells and dnT on D8.

Pro-inflammatory signaling was evident through upregulation of TNF (in CD4⁺ TCM on D8), and cytokines IL32 (in NK and dnT cells on D8), CCL4 (in dnT cells and CD4⁺ TCM on D8), and CCL5 (in dnT cells on D8). These changes are consistent with IFN-driven immune recruitment and further inflammation.

Immunoglobulin-related transcripts, such as IGHV3-72 (in naive B cells on D8) andIGKV1D-17 (in naïve B cells on D8) were upregulated, potentially enhancing antigen recognition and decreasing antibody diversity. T cell activation and cytotoxicity were marked by increased expression of CD69 in CD8⁺ TEM and NK cells on D8, and CD38 in NK cells on D8/10. The cytotoxic granzyme gene GZMA was upregulated in dnT at both post-infection time points, and in naïve CD8⁺cells on D8. In addition, LAMP3 was upregulated in naïve CD4⁺ and CD8⁺ T cells, CD8⁺ TEM, dnT, and on D8 and on D10.

Expression of LAG3, a checkpoint inhibitor that dampens T cell activation and proliferation, was elevated in γδ T cells on D10 and in γδ T and dnT cells on D8. The latter suggests early features of T cell exhaustion. Meanwhile, downregulation of several TCR variable genes (TRBV4-1, TRBV24-1, and TRBV16) in γδ T, and CD8⁺ naïve T cells suggests possible selective TCR repertoire reshaping. Genes associated with cell death, like AIF1, FAS, and GADD45B, were upregulated on both D8 and D10, indicating ongoing cell death and tissue remodeling processes. A modest yet statistically significant reduction in VEGFB in γδ T, and CD8⁺ naïve T cells was observed on D8, suggesting a transient impact on vascular or metabolic pathways.

Finally, a broad up-and downregulation of long non-coding RNAs (lncRNAs) was seen across immune subsets. On D8, this included transcripts in NK cells, dnT, CD8⁺ TEM, naïve CD4⁺, and naïve B cells, while on D10, LINC01163 and LINC00987 were decreased in NK cells.

### Differential surface protein expression analysis

On D8 and D10, respectively 23 and 15 cell surface proteins were upregulated, while 3 and 7 surface proteins were downregulated, across 12 (D10) to 17 (D8) immune cell populations compared to baseline (|log2 FC| ≥ 1; padj < 0.05). Between D8 and D10, 5 surface proteins were upregulated in 5 cell subtypes whereas 14 were downregulated in 11 cell subtypes (|log2 FC| ≥ 1; padj < 0.05). (**Figure 4; Supplementary Table S4**).

**Figure 4.**
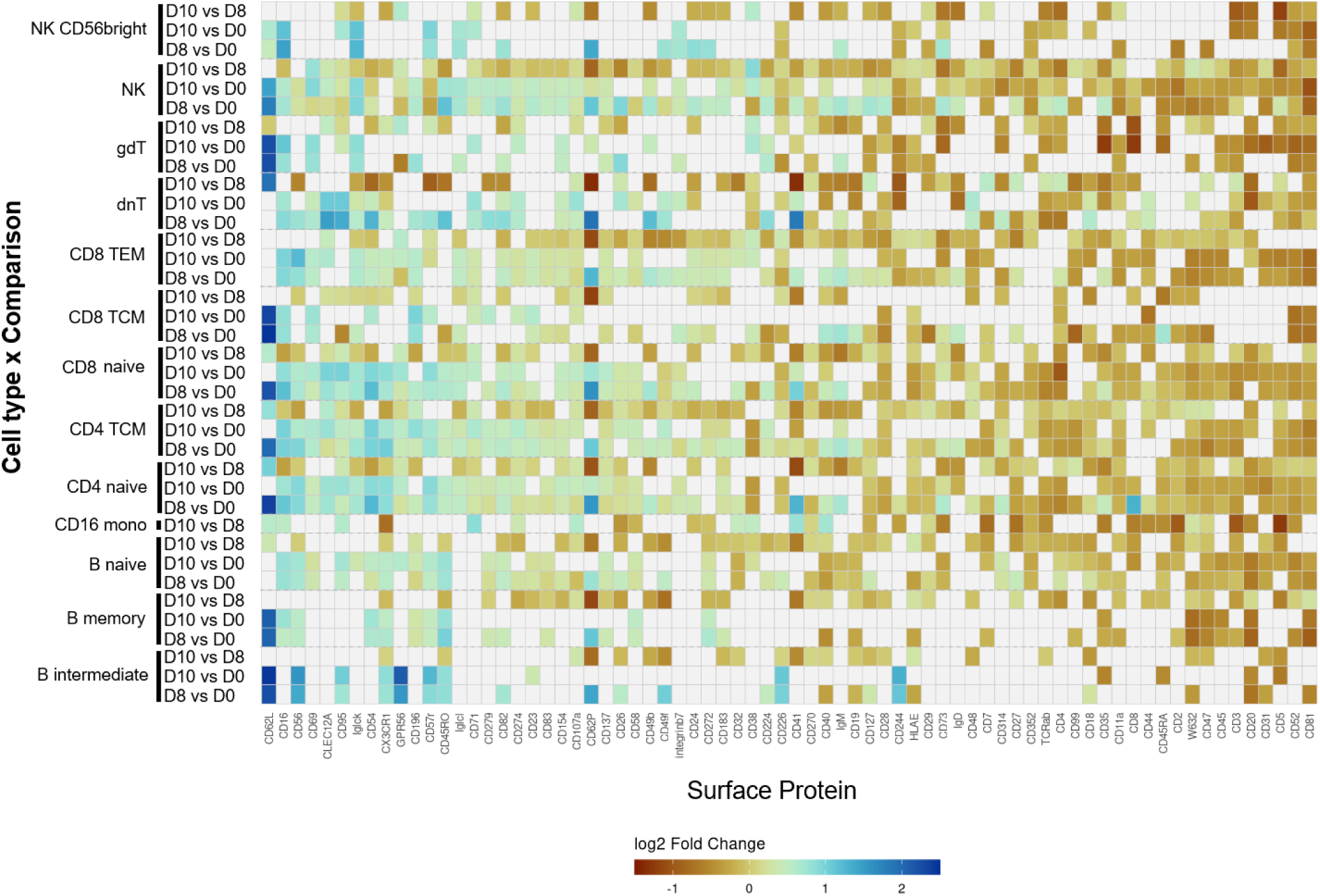
Heatmap of differentially expressed surface proteins. Results were filtered for adjusted p-value < 0.05 and absolute log₂ fold change |logFC| ≥ 1. The x-axis shows surface protein markers and the y-axis lists cell subtypes across time point comparisons. Non-significant values are shown in grey, and cell types without significant differential protein expression or with < 10 different significantly expressed proteins were omitted from the figure.

The adhesion molecule CD62L showed the most widespread upregulation, being significantly increased in major T-cell subsets (CD4⁺ and CD8⁺ naïve, TEM, TCM), γδ T cells, dnT cells, Tregs, NK cells, B cells, and platelets on both D8 and D10. Other migration-related molecules like CD54 (ICAM1) were likewise upregulated on D8 in dnT and naïve CD4⁺ and CD8⁺ T cells, while CD49a also increased in NK cells, and CD41, in dnT cells and CD4⁺ and CD8⁺ T cells, indicating enhanced adhesion and migratory potential. The chemokine receptor CD194 (CCR4) was specifically elevated in dnT cells, supporting an activation and trafficking response.

Several activation and effector-associated receptors were significantly induced, including CD16 (FcγRIII) in B intermediate, NKCD56bright cells, CD4⁺ CTL and TCM, and naïve CD4⁺ and CD8⁺ cells on D8; CD169 in CD16⁺ monocytes (between D8 and D10); CD56 (NCAM) in B intermediate cells on D8/10; and CD101 and CD69 in Tregs and MAIT cells, respectively. Upregulation of GPR56, and KLRG1, and C-type lectin-like receptor 12A (CLEC12A) in respectively intermediate B and dnT subsets further pointed to heightened activation, cytotoxic, and myeloid-like responses.

Conversely, several proteins were significantly downregulated at D10 compared with earlier time points D8 and D0, including CD21 in γδ T cells and proliferating NK cells, CD268 (BAFFR) and CD73 (NT5E) in proliferating NK cells, and TCR Vδ2 in dnT cells. The CD8 protein itself was also decreased in γδ T cells and CD4⁺ naïve cells. These reductions suggest a contraction of naïve or regulatory signaling programs concurrent with effector activation.

The data reveal a coordinated temporal transition marked by strong upregulation of adhesion and activation molecules (CD62L, CD54, CD62P, CD16, CD56) and selective downregulation of receptors associated with naïve or inhibitory states (CD21, CD268, CD73, TCR Vδ2), reflecting immune activation and redistribution between days 8 and 10.

### Gene enrichment analysis

Gene Ontology (GO) analysis identified 390 and 344 significant terms on days 8 and 10 post-challenge, respectively (**Supplementary File S3a; Supplementary Figure S3; Supplementary Figures S5**), alongside 529 and 235 significantly downregulated terms (padj ≤ 0.05). Antiviral defense responses dominated across PBMC subsets, with enrichment of core processes such as viral process and defense response to virus in B and T cell subsets at D8, expanding by D10 to NK, NK CD56bright, and pDC populations, reflecting broader and sustained immune engagement over time.

Interferon-associated categories were similarly prominent and expanded over time. At D8, terms such as “IFN-mediated signaling pathway,” “positive and negative regulation of type I IFN production,” and “type I IFN production” were enriched (padj ≤ 0.05) in B, T and NK cell subsets and persisted at D10, with broader involvement of CD4^+^CTL, CD4^+^TEM, proliferating NK cells, pDCs and Tregs. Consistently, the RIG-I-like receptor signaling pathway, which mediates intracellular viral RNA recognition and IFN-I production, was activated across multiple T-cell subsets at both time points and expanded by D10 to include NK cells, B cells, and Tregs. By D10, this activation extended to include NK cells, B cells, Tregs and CD8⁺ TCM and TEM, reflecting a progressive engagement of innate antiviral sensing. Chemokine signaling and cell migration pathways were particularly enriched on D10. Chemokine signaling and cell migration pathways became more prominent at D10, involving multiple T-cell subsets, B cells, pDCs, and NK populations.

Antigen processing and presentation pathways remained active throughout, enriched in naïve B cells, several T cell subsets and NK cells at D8, and at D10 including pDC. Receptor-mediated sensing pathways such as Toll-like receptor signaling showed a similar expansion pattern, consistent with sustained pattern-recognition receptor activation. Cell death programs were enriched in selected B– and T-cell subsets at D8 and broadened by D10, potentially reflecting the onset of immune contraction. Downregulated GO terms were primarily related to metabolism, gene regulation, development, cytoskeletal and cell cycle processes.

KEGG pathway analysis identified 21 and 20 significantly enriched pathways at D8 and D10, with 10 and 4 pathways significantly downregulated (**Supplementary File S3b; Supplementary Figure S4; Supplementary Figure S6**). Enriched pathways were dominated by mitochondrial dysfunction and proteostasis stress. In addition, microtubule-based intracellular transport pathways were represented, suggesting coordinated metabolic and cellular stress responses. The IFN–RIPK1/3 signaling pathway was enriched in naïve and TCM CD8⁺ cells at D8, expanding to dnT and NK cells by D10. Cytokine JAK–STAT signaling was enriched across multiple T-cell subsets at both time points, whereas the IL-2 family to JAK–STAT signaling pathway was uniquely enriched in intermediate B cells at D10.

Overall, pathway analysis reveals a coordinated antiviral response marked by sustained IFN signaling, broad immune activation, and metabolic stress responses that expand across immune cell subsets from D8 to D10 post-challenge.

### Intercellular interactions between immune cells

To characterize immune cell crosstalk during DENV-1 infection, we inferred ligand–receptor signaling using CellChat (**Supplementary File S4a/b/c/**). Across time points, the number of highly variable ligand-receptor pairs available for analysis was 1,907 on D0, 2,402 on D8, and 2,315 on D10. Communication could not be assessed for several rare or low-viability populations, which were excluded due to insufficient cell numbers: On D0 plasmablasts, CD16^+^ monocytes, HSPCs, proliferating NK, pDCs; On D8 erythroid cells, proliferating CD8^+^ T cells, HSPCs, pDCs; On D10 proliferating CD8^+^ T cells, ILC, plasmablasts, pDCs, HSPCs. The identity of the main sending populations shifted markedly over time. On D0, naïve B cells accounted for the highest number of inferred ligand-receptor interactions (n = 285), primarily T cell subsets (CD8⁺ and CD4⁺ T cells) and NK cell subsets. By D8, CD14⁺ monocytes became the dominant signalers (n = 371), particularly interacting with CD14⁺ and CD16⁺ monocytes, indicating intra-myeloid communication during early inflammation. By D10, the signaling axis consolidated around CD16⁺ monocytes (n = 346). This transition from CD14⁺ to CD16⁺ monocyte-centered signaling likely reflects early innate activation followed by a more specialized effector response, consistent with known roles of CD16⁺ monocytes in antiviral defense and antigen presentation.

Network topology analysis showed that naïve, TCM and TEM CD8⁺ T cell subsets consistently acted as both prominent senders and receivers throughout infection (**Figure 5**). Their dense intralineage connectivity intensified and consolidate by D10, suggesting progressive coordination among cytotoxic T cell compartments. At D0 and D8, signaling among CD8⁺ subsets was mediated primarily by CD8A/B–HLA-A/B/C/E/F interactions, suggesting regulation of cytotoxic activity, potentially through modulation of NK and CD8^+^ T cell responses. Also, CD16^+^ monocytes increased signaling toward naïve, TCM and TEM CD8^+^ T cells and NK cells from D8 to D10, supporting a model in which monocyte-derived cues shape the expansion, activation, and eventual contraction of cytotoxic populations.

**Figure 5.**
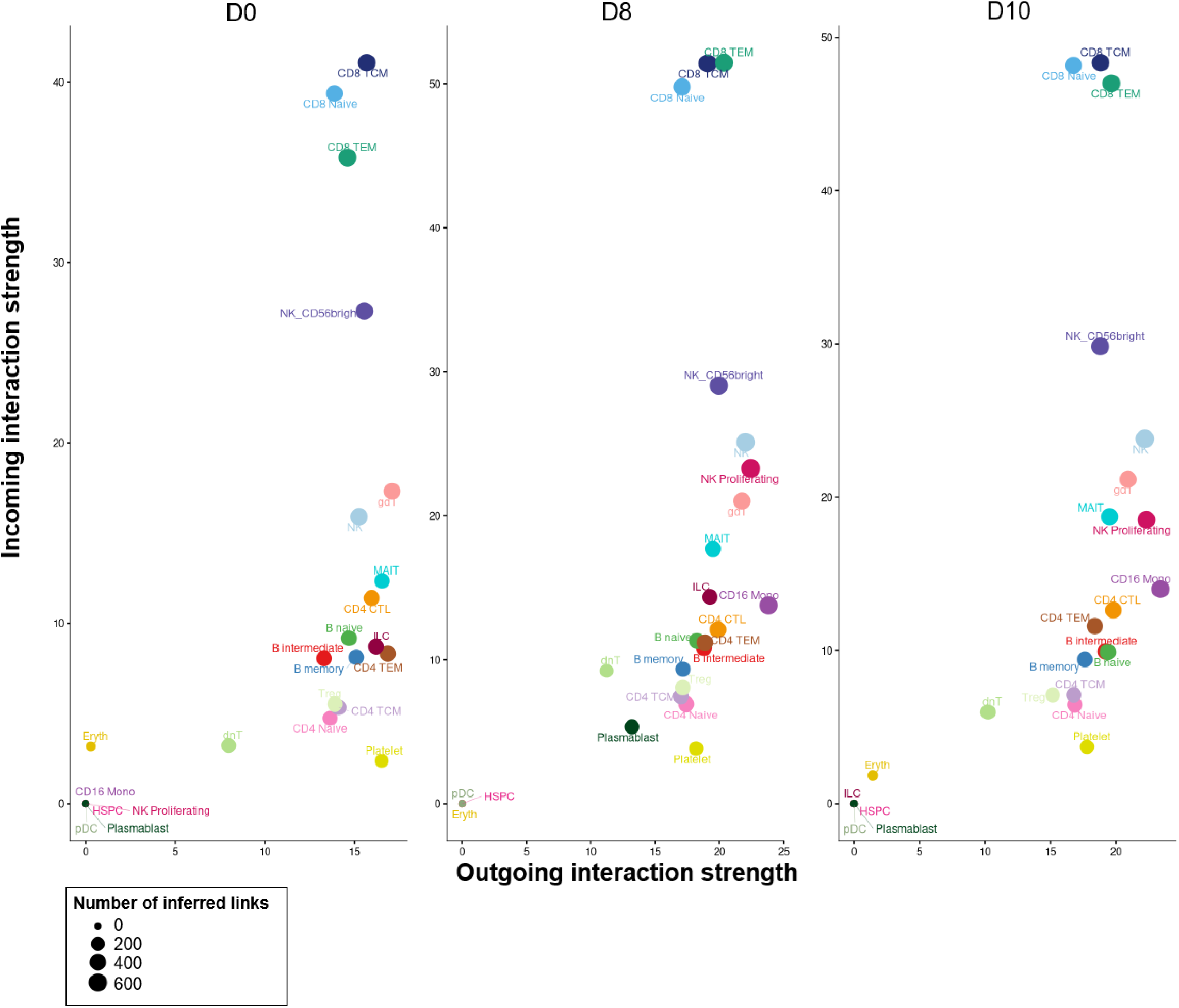
Identification of cell populations with significant changes in intercellular communication by comparing outgoing (ligand) and incoming (receptor) interaction strengths across days 0, 8 and 10. The x-axis and y-axis represent outgoing or incoming communication probabilities for each cell group, respectively. Dot size is proportional to the number of outgoing and incoming interactions associated with each cell type.

Key immunoregulatory interactions were also detected. The MIF-CD74-CXCR4 axis was active at D8 across a broad range of immune populations, including B cells, Tregs, NK cells, MAIT cells, γδ T cells, and multiple CD4⁺ and CD8⁺ T-cell subsets. MIF-CD74 signaling is known to amplify inflammatory recruitment and survival signaling, suggesting that this pathway may help sustain early lymphoid activation during acute infection.

Several additional signaling routes showed cell type and time-specificity. Galectin-9 (LGALS9) signaling through CD44/CD45 was enriched on D8/D10, linking monocytes with T cell and NK cell subsets. Galectin-9 is associated with immune regulation, T-cell exhaustion, and attenuation of overactive responses, and its induction in our dataset likely reflects an early attempt to restrain immune-mediated pathology during primary infection. A similar prominent interaction on D8 involved CLEC2B/D-KLRB1, although DGE analysis revealed KLRB1 expression was restricted to dnT, with a concurrent downregulation of CLEC2D in NK cells. This mismatch between ligand and receptor expression suggests that the CLEC2B/D–KLRB1 pathway may be functionally constrained or suppressed during acute infection, despite computational evidence of signaling potential.

### Viral detection

Dengue virus was not detected in our dataset using either Vulture or CellRanger. However, both pipelines were able to detect DENV-1 in the dataset from Sanborn et al. (2020) (unpublished data).

## Discussion

### Overview and key advances

This longitudinal single-cell multi-omics analysis of experimentally induced DENV-1 infection provides a detailed view of the cellular and molecular dynamics that characterize acute primary dengue infection. While previous single-cell dengue studies have described transcriptional activation, our combined transcriptomic and surface-proteomic profiling, paired with ligand-receptor inference, reveals previously unresolved system-level features of dengue immunity.

In particular, we identify: (1) a monocyte-CD8⁺ T-cell signaling axis that dominates early antiviral coordination, (2) early B cell priming coupled with extensive adhesion and activation remodeling across lymphocytes, and (3) cell type-specific translational and regulatory control shaping antiviral effector activity. These findings refine and extend earlier single-modality studies, which lacked multimodal integration or communication-level resolution, and they provide a mechanistic view of how coordinated antiviral immunity is orchestrated during primary dengue infection.

### A myeloid–cytotoxic signaling hub

A defining feature of this immune landscape was the emergence of a CD16⁺ monocyte-centered signaling architecture (18, 20), with CD8⁺ T cells and NK cells functioning as stable bidirectional partners. This pattern indicates a redistribution of antiviral coordination toward a myeloid-cytotoxic lymphocyte hub, rather than a diffuse innate response. This architecture also contrasts sharply with Zika virus infection, where productive infection of DCs and impaired antigen presentation limits lymphocyte engagement. However, Ghita et al. (2023) reported that antigen-presenting cells retain normal antigen uptake but exhibit impaired IFN signaling and antigen-processing/presentation early in patients who develop SD, alongside a reduced CD16⁺ monocyte fraction (13). In our dataset, the monocyte-centered network developed alongside selective shifts in PBMC composition: dnT cells declined from baseline to D8. This pattern likely reflects a transient IFN-I mediated redistribution or apoptosis rather than permanent depletion (18, 24–27).

Earlier single-cell dengue studies often treated individual cells as independent observations, inflating false discovery rates (28, 29). By applying pseudobulk analysis, we identified more selective and biologically plausible abundance shifts (30). This approach helps reconcile discrepancies across the literature; some studies reported broad B– and T-cell expansion (12, 18), whereas Waickman et al. (2021) observed stable leukocyte proportions before day 14. Such differences likely reflect both methodological choices and the attenuated DENV-1 strain used in the human challenge model, which elicits a milder, more controlled immune response (20).

### Coordinated activation of lymphoid compartments

Cytotoxic and helper T-cell subsets showed upregulation of activation markers including CD69 and GZMA, consistent with effective cytotoxic and helper responses associated with milder dengue outcomes (31, 32). Protein-level measurements uncovered additional remodeling of adhesion and activation receptors, such as ICAM1 and CD62L, across multiple T and B cell subsets, indicating enhanced effector function, homing capacity, and immunoregulatory activity. Interestingly, specific ICAM1 genotypes have been associated with increased susceptibility to dengue, potentially due to altered immune-cell recruitment to sites of infection, further highlighting the relevance of adhesion pathways in disease variability (33). The upregulation of LAG3 in γδ T and dnT cells points to early engagement of regulatory mechanisms that temper effector activity without limiting antiviral function (13). However, unlike Ghita et al. (2023), we did not see typical activation and exhaustion signatures in NK cells (13). B cells likewise exhibited early activation, with induction of immunoglobulin variable region transcripts and surface proteins linked to priming and migration. These multilayered activation signatures extend prior studies by demonstrating that lymphocyte engagement is not merely reactive but structurally coordinated across compartments.

### Interferon-driven antiviral programs

Interferon signaling dominated the immune landscape. Although type I IFNs were not detected, likely due to the lack of very early sampling (34, 35), their downstream targets were strongly induced across cell types. ISGs restrict viral replication and amplify innate immunity, but flaviviruses like dengue and Zika can exploit ISG15-mediated IFN-I regulation to evade innate immunity. Notably, ISG15-deficient humans remain protected against common viral infections such as influenza and HSV-1, likely due to a sustained antiviral IFN-I response (36–38).

IFI27, previously associated with positive regulation of DENV2 replication (12), was highly expressed in CD4^+^ TEM cells, and NK cells on D/810^16^, diverging from early reports placing its expression primarily in monocytes and pDCs, revealing greater ISG heterogeneity than earlier reported^9^. Moreover, broad ISG upregulation confirmed hallmark dengue signatures(12, 20). In contrast, IFNγ expression was enriched in NK cells and CD8^+^ TEM cells, consistent with its peak at defervescence (35). Engagement of RIG-I receptor pathways across lymphoid compartments further highlights the distributed nature of pathogen sensing rather than strictly myeloid-restricted detection, enabling antiviral immunity while limiting excessive inflammation (39, 40). Coordinated upregulation of IRF7 and antigen-presenting signatures supports a model in which IFN programs not only restricts viral replication but also primes adaptive responses through enhanced communication and antigen processing (41, 42). Unlike Zika, which suppresses DC-mediated priming, primary dengue sustains monocyte-T cell coordination and early B-cell priming (43). Although IFN-driven activation is also seen in COVID-19, dengue differs substantially in cytokine magnitude, temporal structure, endothelial involvement, and long-term sequelae (44).

### Translational control

Our multi-omic data also point to translational control as a central regulatory mechanism. Downregulation of EIF4E3 and simultaneous upregulation of EIF2AK2 and EIF5A in NK and T-cell subsets suggest coordinated induction of stress-responsive translational checkpoints shaping antiviral protein synthesis while preserving cellular viability (45). These findings align with viral strategies to manipulate host translation to evade immune restriction while ensuring the synthesis of essential viral and host factors (45). Unlike Waickman et al. (2021), we did not observe repression of mitochondrial genes like MT-CYB, likely due to stringent quality control filtering (20). Together, these translational checkpoints and IFN-driven programs form a core determinant of acute dengue immunity that supports strong early immunity without triggering broad immunopathology.

### Cell death and pathway activation

Increased expression of pro-apoptotic ISGs (46, 47) in T and NK cells indicates activation of cell stress and death pathways, yet concurrent induction of the anti-apoptotic ISG IFI6 suggests a dynamic equilibrium in which cells engage antiviral effector pathways while preventing excessive loss of key immune populations (20, 48). Upregulation of the C-type lectin CLEC12A in dnT cells suggests engagement of negative feedback mechanisms. Although CLEC12 is a principal DENV receptor, its functional role in dengue remains unclear (49, 50). Evidence from LCMV, influenza (51), and picornavirus (52) infections shows that CLEC12A modulates antiviral immunity and T-cell activation, implying similar key regulatory roles in dengue infection.

Single-cell multi-omics enabled unprecedented, detailed mapping of intercellular communication networks. LGALS9-CD44/CD45 interactions between monocytes and lymphoid cells were enhanced, consistent with natural dengue infections (12). Although elevated LGALS9 is associated with severe disease (53), our cohort did not exhibit a cytokine profile characteristic of severe dengue, such as increased IL-2 and IL-6 (10, 11).

### Viral detection

Viral RNA was not detected in any immune subset. This aligns with previous analyses of this human challenge model (11, 54) and reflects low viremia observed in this cohort, rapid turnover of infected cells, and technical limitations of single-cell detection (55, 56), rather than true absence of infection. Variation in peak viremia timing across participants may also have shaped compositional and transcriptional differences. Nonetheless, host transcriptional programs clearly demonstrated antiviral sensing.

### Limitations

Although the cohort size was modest, the controlled nature of a human challenge model minimizes confounding by prior flavivirus exposure, variable infection timing, and heterogenous disease trajectories. The small sample size limits statistical power and generalizability, and the absence of SD cases precludes direct assessment of immunopathogenic mechanisms underlying clinical deterioration. Several samples could not be analyzed due to low viability, likely reflecting dengue-associated leukopenia (57) and the fragility of activated or infected PBMCs, which are susceptible to cryopreservation (58) and virus-induced cytotoxic stress (59). These factors, combined with the rigid viability requirements of the 10x Genomics platform, and cryopreservation artifacts selectively affecting specific PBMC subsets (58, 60), may bias abundance-based analyses. Notably, plasmablasts were not detected in our dataset, representing a missing dimension of the humoral response. Contrastingly, progression to SD has been associated with pronounced plasmablast expansion, highlighting a key divergence between controlled, milder infection and naturally occurring severe disease (13). Moreover, changes in immunoglobulin transcript abundance were revealed, though evaluation of gene usage and clonal diversity would require dedicated V(D)J sequencing.

## Conclusion

In conclusion, this study provides the first integrative single-cell transcriptomic and surface proteomic analysis of acute primary DENV-1 infection. By combining high-resolution profiling of cellular composition, activation states, and intercellular communication, we confirm established dengue signatures such as CD16⁺ monocyte expansion and strong ISG induction, and we uncover additional features including coordinated adhesion remodeling, early B cell activation, cytotoxic T cell engagement, cell-type specific translational control, and myeloid-centered network reorganization. Intercellular communication analysis revealed dense signaling between CD8⁺ T cells, NK cells and monocytes, supporting a coordinated antiviral response. This architecture expands previous models of dengue by clarifying how sensing, coordination, and regulation are distributed across PBMC subsets during primary infection. Together, these findings refine mechanistic understanding of dengue immunopathogenesis, and offer a valuable foundation for biomarker discovery and immune-targeted intervention strategies.

## Data Sharing Statement

Clinical participant data has been previously published(19). Raw CellRanger data are available through the NCBI Gene Expression Omnibus under accession number (GSE314581).

## Contributors

LATW and SJT designed the clinical study. ASV, FVN, KKA and OL designed, analyzed, and interpreted the study. EDM conducted laboratory work associated with the scRNAseq. ASV prepared and finalized the draft manuscript. All authors read and approved the final version of the manuscript.

## Acknowledgements

This work was funded by the Flemish agency for innovation and entrepreneurship (VLAIO; grant HBC.2022.0988) and Johson & Johnson. Funders were not involved in the study design, data collection, data analyses, interpretation, or writing of the manuscript.

We thank J&J Biobank Beerse for logistical support, the J&J dengue compound development team for programmatic support and the study participants for making this research possible. We acknowledge EDM, SD, and SDK of NXTGNT for their technical expertise in performing the experimental procedures and ensuring consistently high-quality sample processing.

